# Resting-state fMRI reveals altered functional connectivity associated with resilience and susceptibility to chronic social defeat stress in mouse brain

**DOI:** 10.1101/2024.03.28.587210

**Authors:** Derek Lupinsky, Md Taufiq Nasseef, Carine Parent, Kelly Craig, Josie Diorio, Tie-Yuan Zhang, Michael J. Meaney

**Affiliations:** Douglas Hospital Research Centre, Department of Psychiatry, McGill University, Montréal, Canada; Ludmer Centre for Neuroinformatics and Mental Health, McGill University, Montréal, Canada; Department of Mathematics, College of Science and Humanity Studies, Prince Sattam Bin Abdulaziz University, Saudi Arabia; Translational Neuroscience Program, Singapore Institute for Clinical Sciences, Agency for Science, Technology and Research (A*STAR), Singapore; Yong Loo Lin School of Medicine, National University of Singapore, Singapore

**Keywords:** resting-state, fMRI, resilience, functional connectivity, social defeat, mouse

## Abstract

Chronic stress is a causal antecedent condition for major depressive disorder and associates with altered patterns of neural connectivity. There are nevertheless important individual differences in susceptibility to chronic stress. How stress-induced alterations in functional connectivity amongst depression-related brain regions associates with resilience and susceptibility to chronic stress is largely unknown. We used resting-state functional magnetic resonance imaging (rs-fMRI) to examine functional connectivity between established depression-related regions in susceptible (SUS) and resilient (RES) adult mice following chronic social defeat stress (CSDS). Seed-seed FC analysis revealed that the ventral dentate gyrus (vDG) exhibited the greatest number of group differences in functional connectivity with targeted brain regions. SUS mice showed greater functional connectivity between the vDG and subcortical regions compared to both control (CON) or RES groups. Whole brain vDG seed-voxel analysis supported seed-seed findings in SUS mice and indicated significantly decreased connectivity between the vDG and anterior cingulate area compared to CON mice. Interestingly, RES mice exhibited enhanced connectivity between the vDG and anterior cingulate area compared to SUS mice. Moreover, RES mice showed greater connectivity between the infralimbic prefrontal cortex and the nucleus accumbens shell. These findings indicate unique differences in functional connectivity patterns in SUS and RES mice that could represent a neurobiological basis for vulnerability for stress-induced depression.

## INTRODUCTION

Major depressive disorder (MDD) is a debilitating psychiatric disorder projected to become the main disease burden by 2030 and is now the leading cause of years lived with disability (YLD) (1). Approximately half of depressed patients do not reach a full recovery and relapse is common following treatment (2–5). Chronic stress is often a precipitating risk factor in the emergence of depression (6–9). However, important individual differences exist in the susceptibility for depression following exposure to chronic stress (3, 10–15). The identity of the mechanisms underlying such variation in susceptibility and resilience in response to chronic stress is therefore of seminal importance for an understanding of the pathophysiology of depression. Analyses of relevant rodent models reveal specific transcriptomic profiles associated both resilience and susceptibility to chronic stress (16, 17). Importantly, profiles associated with resilience are distinct from those of both susceptible and control animals. These findings reinforce the perspective that resilience and susceptibility in the face of stress are distinct neural processes. However, we lack a full understanding of the neural circuitry underlying each state.

The chronic social defeat stress (CSDS) model is a highly validated and widely-used rodent model for studies of the mechanisms of stress-induced depression- and anxiety-like behaviors. The CSDS model also reveals variation in susceptibility and resilience. These individual differences are apparent in multiple outcome measures including depressive- and anxiety-like behaviors, metabolic homeostasis and neuroendocrine and immune signals (18, 19). CSDS has thus emerged as an important model for the study of epigenetic, transcriptional, and anatomical mechanisms associated with the pathophysiology of stress-induced depression (13,18, 20–23). A detailed analysis of transcriptional profiles revealed highly significant overlap in gene networks associated with MDD in post-mortem human brain and those resulting from exposure to CSDS in mice (24) attesting to the relevance of the model for studies of the biological basis of MDD.

Depression is a circuit level-wide disorder involving many interconnected brain regions (25, 26) (16, 27–30). Circuit level analysis of activity patterns derived from exposure to CSDS reveal convergence on the ventral hippocampus (31). A previous neuroimaging study with mice following exposure to CSDS showed that structural covariance between the hippocampus and the nucleus accumbens is significantly associated with susceptibility to the effects of CSDS (32). Subsequent analyses identified projections specifically from the dentate gyrus of the ventral hippocampus (vDG) to the nucleus accumbens as a neural circuit that defines susceptibility to CSDS (33). Anacker et al. showed how vDG activity directly mediates individual differences in the susceptibility to stress such that activation of the region produces susceptibility, while silencing vDG excitability through enhanced neurogenesis produces resilience following CSDS. Enhanced glutamatergic tone from ventral hippocampal afferents to the nucleus accumbens enhances susceptibility to CSDS (34, 35). These convergent findings identify the vDG as a critical region for susceptibility to chronic social stress (36, 37).

The hippocampus is a heterogeneous brain structure with the dorsal and ventral portions serving different functions (38, 39) including those related to mood disorders (40). While the rodent dorsal hippocampus is involved in spatial learning/memory, the ventral hippocampus is integral for emotional processing. In addition to the vDG, the mesolimbic dopamine system is implicated in susceptibility to CSDS (41, 42). The basolateral amygdala, a brain region that processes fear- and threat-related stimuli and is likewise implicated in the susceptibility to chronic stress along with other stress-related brain regions; including, the hypothalamus and the periaqueductal gray that modulates behavioural responses to threatening stimuli and pain (32, 43, 44). In contrast, resilience to chronic stress is associated with regions that support effective emotional regulation. The anterior cingulate area is considered a central hub region for the integration of information involved in cognition and decision-making from the prefrontal cortex with information involved in emotionality and emotional learning processed through several areas within the limbic system (45). Accordingly, the anterior cingulate area is critical for ‘top-down’ regulation of emotional activity within the limbic system. Identifying individual differences in connectivity across these brain regions following exposure to chronic stress is a fruitful approach to identifying the neurobiological basis of susceptibility and resilience to chronic stress.

We used resting-state functional magnetic resonance imaging (rs-fMRI) to study brain network changes using functional connectivity (FC) analyses in susceptible and resilient mice following CSDS compared with control, non-stressed mice. Measures of FC are an established approach to study whole brain network connectivity in mice (46). Previous rs-fMRI studies report differences in network communication in response to psychosocial stress (47) and individual variability (48). Our initial analyses used an unbiased, data-driven spatial independent component analysis (ICA) and showed that the vDG, basolateral amygdala, nucleus accumbens, and anterior cingulate area were common components across all groups tested following CSDS. Additionally, seed-seed FC analysis identified the vDG (top aspect) as the region showing the highest number of FC group differences attesting to the role of this brain region as a hub in mediating susceptibility and resilience to CSDS. Based on these findings the vDG (top aspect) was used as a source seed to review whole brain seed-voxel FC. Overall, susceptible (SUS) mice exhibited greater FC between the vDG and other stress-related limbic brain regions. In contrast, in resilient (RES) mice FC between the vDG and prefrontal regions, including the anterior cingulate area, was enhanced compared to SUS mice, implying the possibility of greater top-down cortical regulation of the vDG and perhaps other limbic structures in RES mice. These findings suggest important individual differences in corticolimbic connectivity between SUS and RES mice as the neurobiological basis for the development or resistance to the psychopathology underlying stress-induced depression.

## METHODS

### Animals

C57BL/6 mice were obtained from Charles River Laboratories (Saint-Constant, Quebec, Canada) and bred at the Douglas Research Centre (Montréal, Quebec, Canada). Housing was maintained in a temperature (21±1 °C) and humidity (55±10%) controlled room with a 12 h light: 12 h dark light cycle (lights on 08:00 and 20:00 h). All procedures were performed according to guidelines of the Canadian Council on Animal Care with an animal use protocol (AUP #7239) approved by the McGill University and Douglas Research Centre Facility Animal Care Committees (FACC). Male mice were weaned at postnatal day 22 (PND22) and siblings were assigned to standard housing conditions with mouse chow and water available *ad libitum* for 8 weeks. Animals were raised in groups of three mice (from different litters) in a 30 × 18 cm cage with standard bedding, two nestlet squares, and Enviro-dri available as nesting materials. CD1 retired breeders were obtained from Charles River Laboratories (Saint-Constant, Quebec, Canada) and single-housed.

### Chronic Social Defeat Stress (CSDS)

We followed a published protocol for the CSDS procedures (49). On PND 80 adult male mice were physically defeated for 10 days by a new, pre-screened CD1 aggressor mouse for 5 minutes daily and subsequently housed across a perforated plexiglass divider in the same cage with the aggressor for 24 h. To mimic these conditions, control mice were housed in a cage across a divider with a novel C57BL/6 mouse every day for 10 days. On day 11, animals were screened on the social interaction test. During the first trial, mice were allowed to explore an open field arena (42 cm × 42 cm × 42 cm) containing an empty wire mesh enclosure for 2.5 mins. During the second trial, a CD1 mouse was placed into the enclosure and the experimental mouse was reintroduced for 2.5 mins. Social interaction ratios were calculated as time spent in the interaction zone during the second trial divided by the time spent in the interaction zone during the first trial. Mice with SI ratios <1 were considered “susceptible”; mice with social interaction ratios >1 were considered “resilient”(49). While arbitrary, this approach is consistent with previous studies in the literature to allow direct comparison with existing data sets. Group social interaction scores were assessed for normality using the Kolmogorov-Smirnov test and outlier analysis was performed accordingly using the interquartile range method for non-parametric data or group mean plus or minus two standard deviations for parametric data. All animals were then pair-housed with same group partners in standard cages until brain imaging.

### Resting-state functional magnetic resonance imaging (rs-fMRI)

#### Anesthesia and physiological maintenance

One week following SI testing, mice were weighed (27.98g ± 0.29 SEM) and prepared for rs-fMRI data acquisition. Mice were quickly anesthetized using isoflurane (ISO) (induction: 5% ISO; maintenance: 2% ISO volume mixed with oxygen) for five minutes. Mice were then transferred to the scanner and Cryoprobe bed assembly while ISO anesthesia was maintained between 1.0-1.5% (50). Overall, ISO anesthesia was maintained at an average of 1.29% ± 0.04 SEM. Body temperature during scanning was maintained using a warm airflow over the animal set at 37℃. Respiration rate was monitored using the 1025-IBP-50 Small Animal Monitoring Gating System (SA instruments) and was maintained at an average of 91.2 ± 1.5 (breaths per minute ± SEM).

#### rs-fMRI Image Acquisition

Data acquisition was achieved using a 7T small animal scanner (BioSpec 70/30USR, Bruker) equipped with a Cryoprobe. All fMRI images were obtained following Spin Echo EPI pulse sequence with parameters: FOV: 3.5 × 1; matrix: 128 × 80; slices: 15; resolution: 139 × 125 × 700 μm3; TE/TR: 1/1.50 s; flip angle 90; repetitions: 400(50).

#### rs-fMRI preprocessing

Preprocessing involved: (i) denoising using Advanced Normalized tools (ANTs, *DenoiseImage*) (github.com/ANTsX/ANTs/blob/master/Examples/DenoiseImage.cxx); (ii) motion correction using ANTs and FSL toolboxes (fsl.fmrib.ox.ac.uk/fsl/fslwiki); (iii) co-registration was performed by using ANTs whereby a functional template was created using all subjects’ mean volumes and to normalize each subject’s data; (iv) bias field correction was applied using ANTs; (v) using MINC tools (www.bic.mni.mcgill.ca/ServicesSoftware) we manually registered subject fMRI data from step iv to a high-resolution Allen Mouse Brain template (http://help.brain-map.org/display/mouseconnectivity/API, annotation/mouse_2011) that was down-sampled for optimal fit, as previously described (50), which effectively achieves an isotropic voxel resolution 150 × 150 × 150 μm3 suitable for further analyses and referencing according to labelled regions of the Allen Mouse Brain Atlas (2011); (vi) ventricle signal regression performed by generating a CSF mask and regressing out these signals using Analysis of Functional NeuroImages (AFNI) tools; (afni.nimh.nih.gov/); (vii) smoothing with a Gaussian Kernel of full width half maximum (FWHM) at 0.6 by AFNI tools; (viii) cropping using a brain mask generated according to grey and white matter segmentation using the Allen template space and applied to all subjects; (ix) bandpass filtering between 0.01 Hz to 0.1 Hz was applied by AFNI tools.

#### Post-processing

We applied three commonly used methods to investigate rs-FC from rs-fMRI data: (i) low component spatial independent component analysis (ICA-20), (ii) seed-seed analysis, and (iii) whole brain seed-voxel analysis.

*Independent Component Analysis (ICA).* Data-driven ICA was performed separately in SUS, RES, and CON mouse groups using GIFT tools (Group ICA of fMRI Toolbox, v1.3i, www.nitrc.org/projects/gift/) in MATLAB. We selected 20 components to investigate both larger brain networks and regional activity. ICASSO was used to examine the stability and reliability of the components by randomization and bootstrapping. Components among all three mouse groups were reviewed to identify common stress-related brain regions.

*Seed-seed analysis* was performed considering *a priori* interests of previously established stress-related brain regions. We selected 14 regions including, the vDG—top and bottom aspects, medial prefrontal cortex subregions including the anterior cingulate, prelimbic, and infralimbic cortex, nucleus accumbens shell, bed nucleus of the stria terminalis, habenula, basolateral and central amygdalar nuclei, ventral tegmental area, periaqueductal gray, dorsal raphe nuclei, and the locus coeruleus. Bilateral/central seed masks were modified as needed from the Allen Mouse Brain Atlas (2011). An in-house built MATLAB script(50) was used for computation and statistical analysis. Mean time-series from selected seeds were extracted from each rs-fMRI acquisition for seed-seed correlation analysis and FC statistical testing between selected groups (t-test; threshold T > 2; p= 0.05; FDR corrected).

*Whole brain seed-voxel analysis.* Seed-voxel analysis was performed to investigate brain-wide FC patterns with a vDG (top) seed. Briefly, correlation analysis was performed using FSL tools whereby the mean of the vDG (top) seed time-series was used to produce a whole brain correlation map at the voxel level for each subject, values were then transformed into Fisher Z scores and group comparison analysis was performed using a two-sample t-test (threshold T > 2; p= 0.05). All results were corrected for family-wise error rate (FWER) following clustering with p=0.05(51). Mango tools (http://ric.uthscsa.edu/mango/) were employed to construct 3D correlation maps and for 3D representation of the vDG (top) seed overlaid on the Allen Mouse Brain Atlas (2011) template image (help.brain-map.org/display/mouseconnectivity/ API, annotation/mouse_2011). We used an ‘in-house’ MATLAB script for quantification analysis of seed-voxel results within select brain regions (50). BrainNet Viewer (www.nitrc.org/projects/bnv/) tools were used for 3D representation of findings overlaid on the Allen Mouse Brain Atlas (2011) image. Boxplots represent subject-level correlation coefficients (CC) illustrating FC group differences between the vDG (top) seed and a single (peak) voxel (highest t-stat value) or a common voxel within a specific brain region (i.e. ACA) (Nasseef et al., 2018). R software package (R version 4.1.3, RMINC version 1.5.3.0, pyminc/v0.56, anaconda version 2022.05, minc-toolkit-v2/1.9.18.2, minc-stuffs version v0.1.25, rstudio version 2022.2.3+492, www.r-project.org) was used for plotting.

## RESULTS

### Social interaction tests

The research design is summarized in **Figure 1**, including the CSDS model, MRI data acquisition and processing. Stress-induced alterations in social interactions are commonly used to identify animals susceptible to chronic social defeat stress (18,22,49). Animals characterized on the basis of their social interaction ratio revealed evident clustering in defeated mice such that SUS (0.34 ± 0.05, mean ± SEM; n=20) and RES (2.48 ± 0.40, mean ± SEM; n=6) animals differed significantly in social interaction ratios (t= 9.627 df= 24, p< 0.0001) (**see Supplementary Figure S1**). One-way ANOVA showed a significant effect of the group (F (2,41) =13.21, p<0.0001) and *post hoc* testing showed that SUS mice had a significantly lower social interaction ratio than CON (p<0.0001) and RES mice (p< 0.001) (see Supplementary Figure 1). Inclusion of the data from a marginal outlying CON had no effect on the findings described below.

**Fig. 1.**
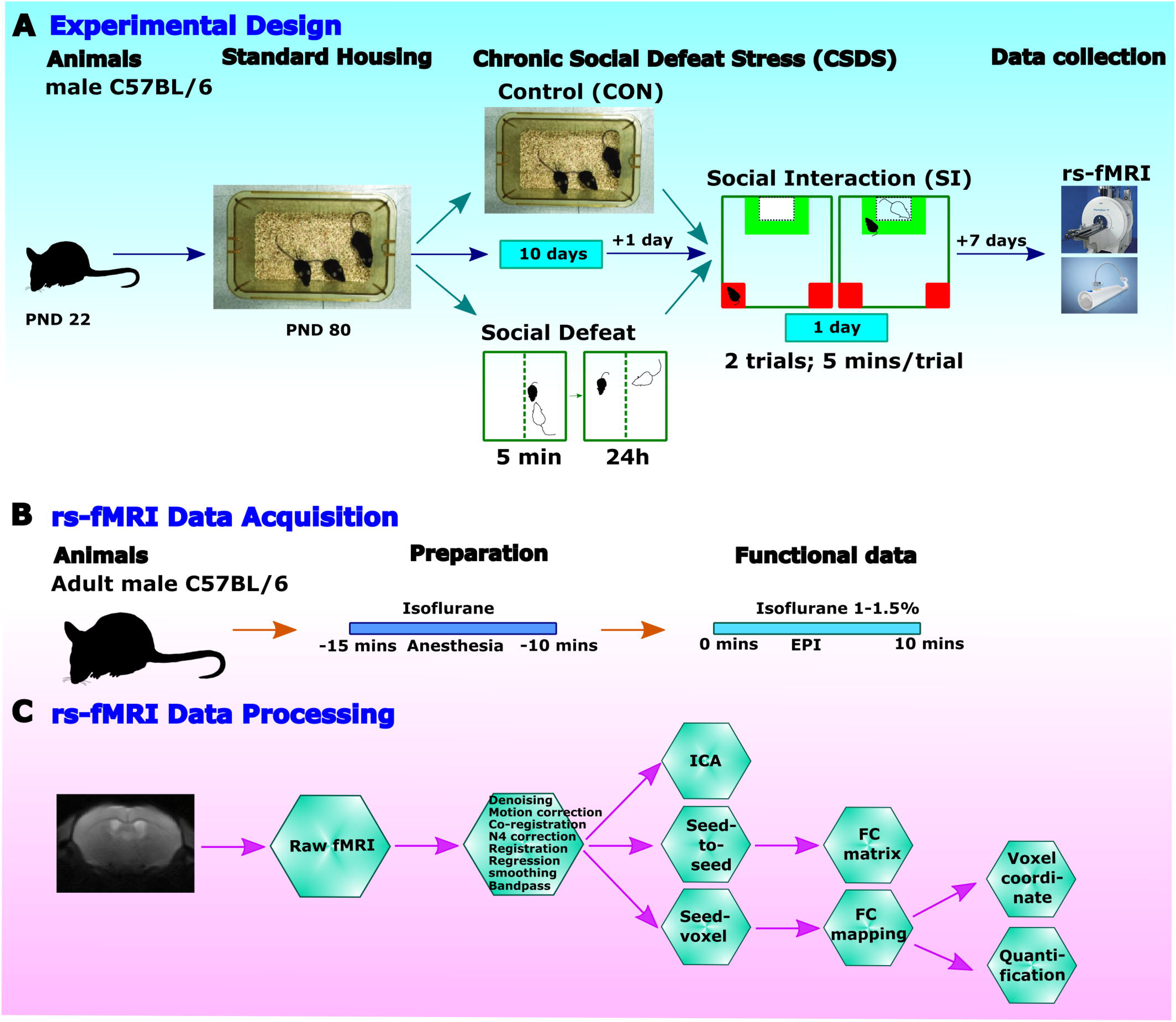
Experimental design, rs-fMRI data collection and processing. A) Experimental design: Male mice weaned on PND22 were assigned to standard housing conditions until PND80 prior to 10 days of chronic social defeat stress (CSDS) or continued standard housing (CON). Social interaction (SI) testing characterized defeated animals as ‘susceptible’ (SUS) or ‘resilient’ (RES) according to their SI ratio. All animals were then pair-housed in standard conditions with appropriate group members for 7 days prior to rs-fMRI. B) A schematic timeline of rs-fMRI acquisition with Cryoprobe technology. C) rs-fMRI data processing and post analysis workflow.

### Spatial independent component analysis (ICA) of rs-fMRI data

Low component spatial ICA was performed separately in SUS, RES and CON mice to identify larger resting-state network components. Components representing established stress-related brain regions including the vDG, basolateral amygdala, nucleus accumbens, and the anterior cingulate cortex that were common to all groups (**Fig. 2A-C**).

**Fig. 2.**
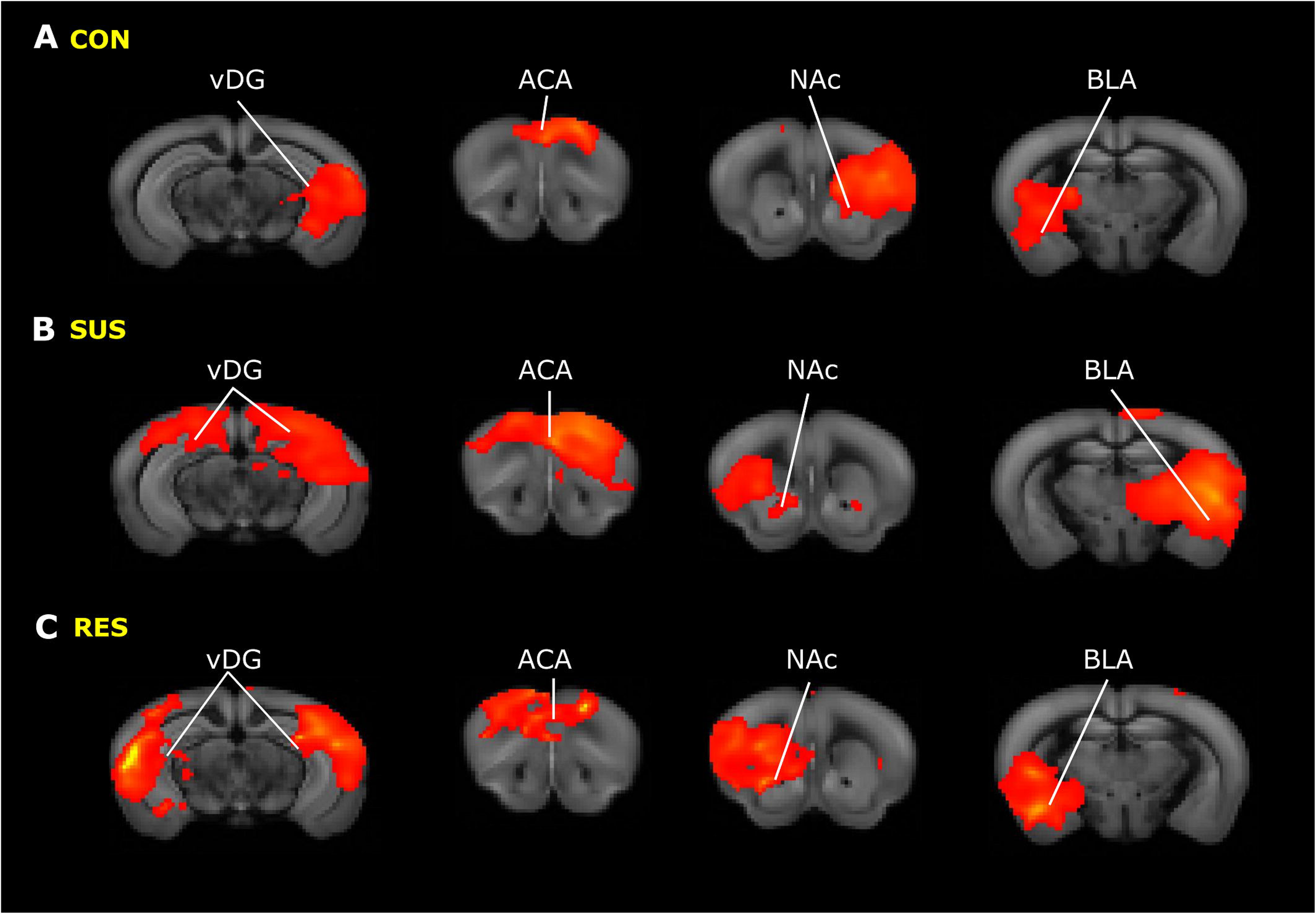
Low component ICA spatial maps in CON, SUS, and RES mouse groups. A-C) Findings from low component (20) spatial ICA performed on all rs-fMRI data from individual mouse groups (A. CON (n=18), B. SUS (n=21), & C. RES (n=6) identified four common stress-related components, specifically the vDG, ACC, NAc, and BLA regions in all animals. Prominent components (red-yellow) are overlaid on corresponding template coronal brain slices in identified mouse groups. *Abv*: ACC, anterior cingulate cortex; BLA, basolateral amygdala; CON, control; NAc, nucleus accumbens; RES, resilient; SUS, susceptible; vDG, ventral dentate gyrus.

### Seed to seed functional connectivity analysis amongst stress-related brain regions

We focused on fourteen stress-related brain regions to conduct hypothesis-driven, seed-seed analysis of rs-fMRI data. Bilateral/central seeds were selected by referring to the Allen Mouse Brain Connectivity Atlas (2011) that included the anterior cingulate cortex, prelimbic and infralimbic prefrontal cortices, nucleus accumbens shell, bed nucleus of the stria terminalis, habenula, basolateral and central amygdalar nuclei, ventral tegmental area, periaqueductal grey, dorsal raphe and locus coeruleus regions and represented on the template atlas 3D image (**Fig. 3A**). Seeds for the vDG (top) and vDG (bottom) regions were manually generated by referring to the Allen Mouse Brain Connectivity Atlas (2011) and represented with other 3D seeds (**Fig. 3A**). Subject-level time-series of each seed region were extracted from rs-fMRI data and correlated with that of all other seeds to compute group mean correlation coefficients (**Fig. 3B**). Significant within group seed-seed FC were evaluated using a one-sample t-test (p<0.05, FDR cut-off 0.05) (**Fig. 3C**). Seed-seed FC group differences were calculated using a two-sample t-test (p<0.05, −2>t>2, FDR cut-off 0.05) that identified significant seed-seed FC differences (**Fig. 3D**).

**Fig. 3.**
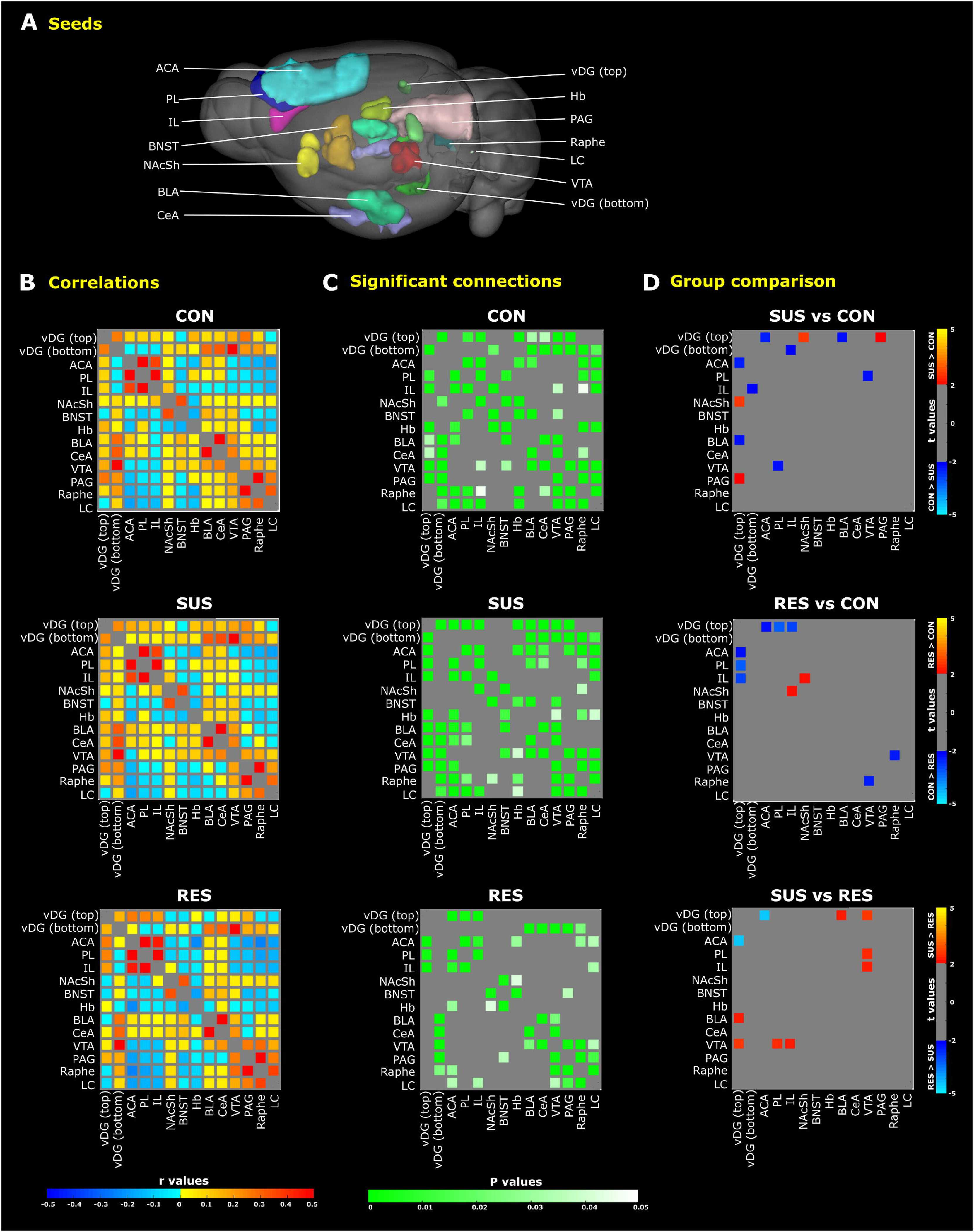
Seed-seed functional connectivity (FC) amongst 14 stress-related brain regions. A) 3D representation of 14 brain regions (seeds) overlaid on a whole brain image of the Allen Mouse Brain Connectivity Atlas (2011). B) Matrices depicting group mean correlation coefficients (*r* values) indicate generally more negative correlations between seed pairs in RES mice (*bottom panel*; blue, cool colours). C) Matrices depicting significant seed-seed connections determined by one sample t-tests (p<0.05, FDR corrected) indicate stronger connectivity (green-white colour scaling) among stress-related brain regions in SUS mice (*middle panel*). D) Statistically significant group differences for a particular seed pair as assessed by two sample t-tests (p<0.05, −2> t > +2, FDR corrected) are indicated. SUS mice feature vDG hyperconnectivity with several subcortical stress-related brain regions (red-yellow colour) when compared to CON (*top panel*) and RES (*bottom panel*) mouse groups. Compared to SUS mice, preferential vDG–ACA connectivity (blue-light blue colour) is a common finding among both RES (*bottom panel*) and undefeated CON mice (*top panel*). *Abv*: ACA, anterior cingulate area; BLA, basolateral amygdala; BNST, bed nucleus stria terminalis; CeA, central nucleus amygdala; Hb, habenula; IL, infralimbic prefrontal cortex; LC, locus coeruleus; NAcSh, nucleus accumbens shell; PAG, periaqueductal gray area; PL, prelimbic prefrontal cortex; RES, resilient; SUS, susceptible; vDG, ventral dentate gyrus; VTA, ventral tegmental area.

The vDG (top) seed showed the greatest number of significant FC group differences among selected brain regions. When compared with CON mice, the SUS but not RES mice showed significantly greater FC (p<0.05, −2 > t > +2, FDR corrected) between the vDG and periaqueductal grey as well as the nucleus accumbens shell region. SUS mice also demonstrated significantly greater FC between the vDG and ventral tegmental area as well as the basolateral amygdala when compared to RES mice (**Fig. 3D**). In contrast, by comparison to SUS mice, the RES mice displayed mostly negative correlations and reduced FC amongst the same brain regions (**Fig. 3D**). The SUS mouse brain is thus best characterized by hyperconnectivity between the vDG and stress-related limbic regions compared to both CON and RES mice. We note that both RES and SUS animals showed reduced connectivity between vDG and ACC, that nevertheless the RES mice showed significantly increased FC between the vDG and anterior cingulate area compared to SUS mice (**Fig. 3D**). RES mice also showed significantly increased FC between the nucleus accumbens shell and infralimbic prefrontal cortex compared to SUS and CON mice (**Fig. 3D**). The RES, by comparison to the SUS mouse brain, is thus best characterized by functional hyperconnectivity between subcortical (vDG or nucleus accumbens shell) and cortical brain regions including the anterior cingulate and infralimbic cortices. Of note there was also significantly increased FC between the ventral tegmental area and the infralimbic and prelimbic subregions of the prefrontal cortex in SUS compared to RES mice (**Fig. 3D**). Further rs-FC analyses focused on the vDG (top) seed since this region showed the greatest number of significant differences compared with other brain regions.

### Whole brain vDG seed-voxel functional connectivity analysis in SUS and RES mouse brain

We then extended our analyses to review brain-wide FC with a vDG (top) seed (**Fig. 4A**) at the voxel level. Group differences of vDG (top) seed-voxel FC identified preferential connectivity patterns specific to individual groups using two sample t-tests (−2>t>2, p<0.05, FWER corrected 0.05) (**Fig. 4B-D**). We again found evidence for hyperconnectivity between stress-related subcortical regions in the SUS mouse brain using seed-voxel analysis revealing convergence across the analytical approaches. SUS mice showed significantly increased FC between the vDG (top) and the nucleus accumbens compared to CON mice (**Fig. 4B**). SUS mice also showed significantly increased FC between the vDG (top) and the basolateral amygdala, hypothalamus, ventral tegmental area, and periaqueductal grey regions compared to RES mice (**Fig. 4D**).

**Fig. 4.**
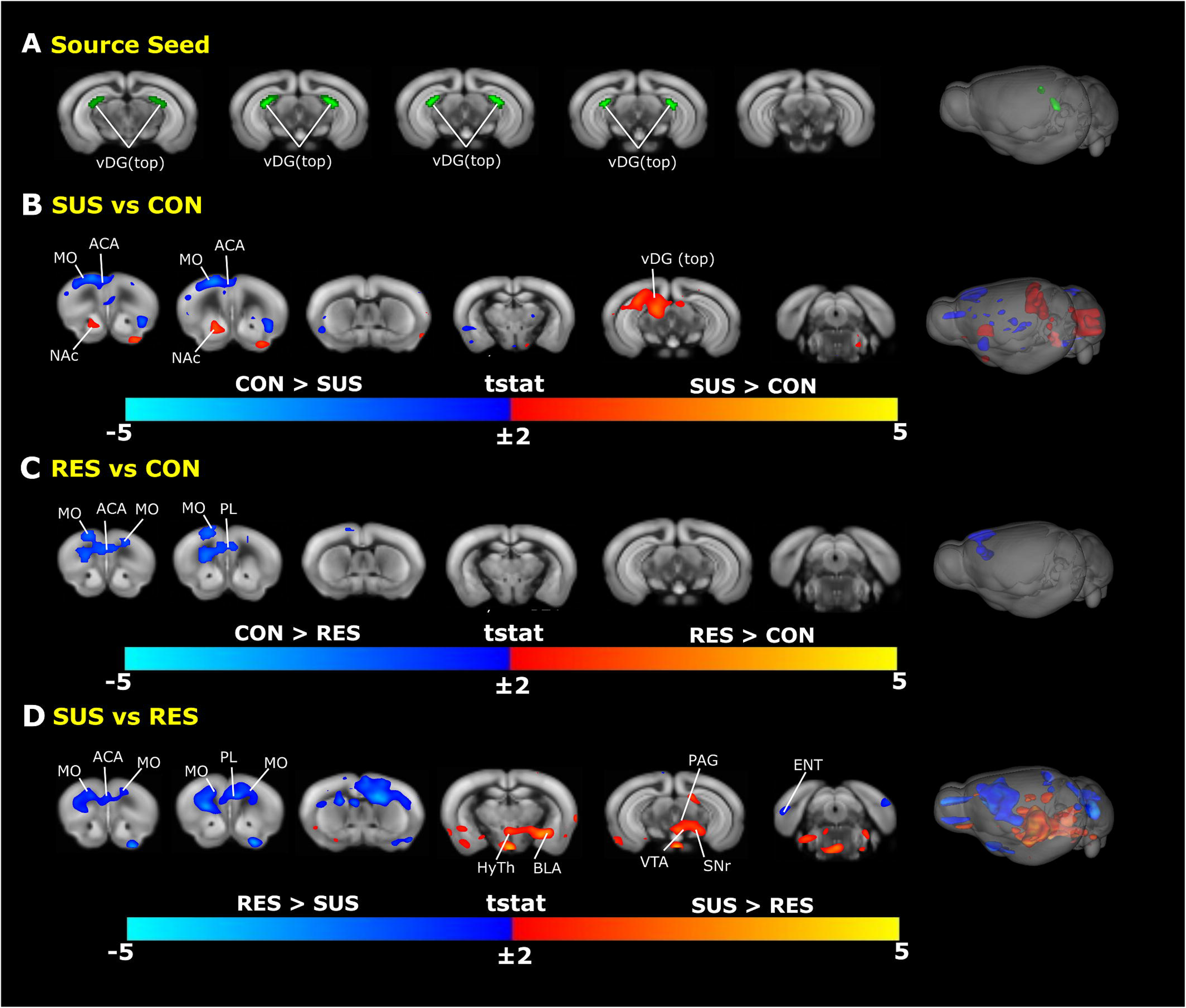
Whole brain vDG (top) seed-voxel functional connectivity (FC) analysis. A) Representations of a manually-generated vDG (top) seed overlaid on Allen Mouse Brain Connectivity Atlas (2011) 2D coronal slices and 3D whole brain image. B-D) Significant group differences of vDG (top) seed-voxel FC are depicted as group contrast t-stat maps overlaid on Allen Mouse Brain Connectivity Atlas (2011) 2D coronal slices and 3D whole brain image; red-yellow (t>2, cc=0.05, FWER-corrected) and blue-light-blue (t<-2, cc=0.05, FWER-corrected) colour intensity. SUS mice display vDG hyperconnectivity with the NAc region when compared to CON mice (B) as well as with the BLA, HyTh, VTA and PAG regions when compared to RES mice (D). Preferential connectivity between the vDG and prefrontal cortical regions (blue-light blue colour) differentiates undefeated CON mice (B) and RES mice (D) from SUS mice. *Abv*: ACA, anterior cingulate area; BLA, basolateral amygdala; BNST, bed nucleus stria terminalis; ENT, entorhinal area; HyTh, hypothalamus; MO, motor cortex; NAc, nucleus accumbens; PAG, periaqueductal gray area; PL, prelimbic prefrontal cortex; RES, resilient; SNr, substantia nigra; SUS, susceptible; vDG, ventral dentate gyrus; VTA, ventral tegmental area; CON, control.

We also found differences in FC between the vDG (top) seed and cortical regions. Importantly, RES mice again showed significantly stronger FC between the vDG (top) and anterior cingulate area as well as the prelimbic prefrontal cortex compared to SUS mice (**Fig. 4D**). Seed-voxel findings were overall supportive of the seed-seed analyses and further demonstrate that the RES mouse brain is best characterized by functional hyperconnectivity between the vDG (top) and cortical regions.

### Functional connectivity contrasts between the vDG seed and peak and common ACC voxel

A common finding amongst seed-voxel group contrasts performed above (**Fig. 4B-D**) was that of significant differences in FC between the vDG (top) and anterior cingulate cortex. We sought to further evaluate these differences by considering connectivity between the vDG (top) seed and the peak voxel (highest t-value) within the anterior cingulate cluster from each group contrast. We found that the location and magnitude of the peak anterior cingulate area voxel depended on the specific group contrast (**Fig. 5A-C top panels**). The peak anterior cingulate area voxel for the SUS versus RES group contrast was localized more posterior within this region with significantly reduced connectivity with the vDG in the SUS group (**Fig. 5C top panel**). RES mice showed significantly increased FC between the vDG (top) and anterior cingulate area peak voxel when compared to SUS mice (t(25)=4.14, p=0.0003, FDR cut-off 0.05) with coordinate (44,67,36) (**Fig. 5C bottom panel**). These results further support functional hyperconnectivity between the vDG and anterior cingulate regions as a possible biological mechanism for resilience following chronic stress.

**Fig. 5.**
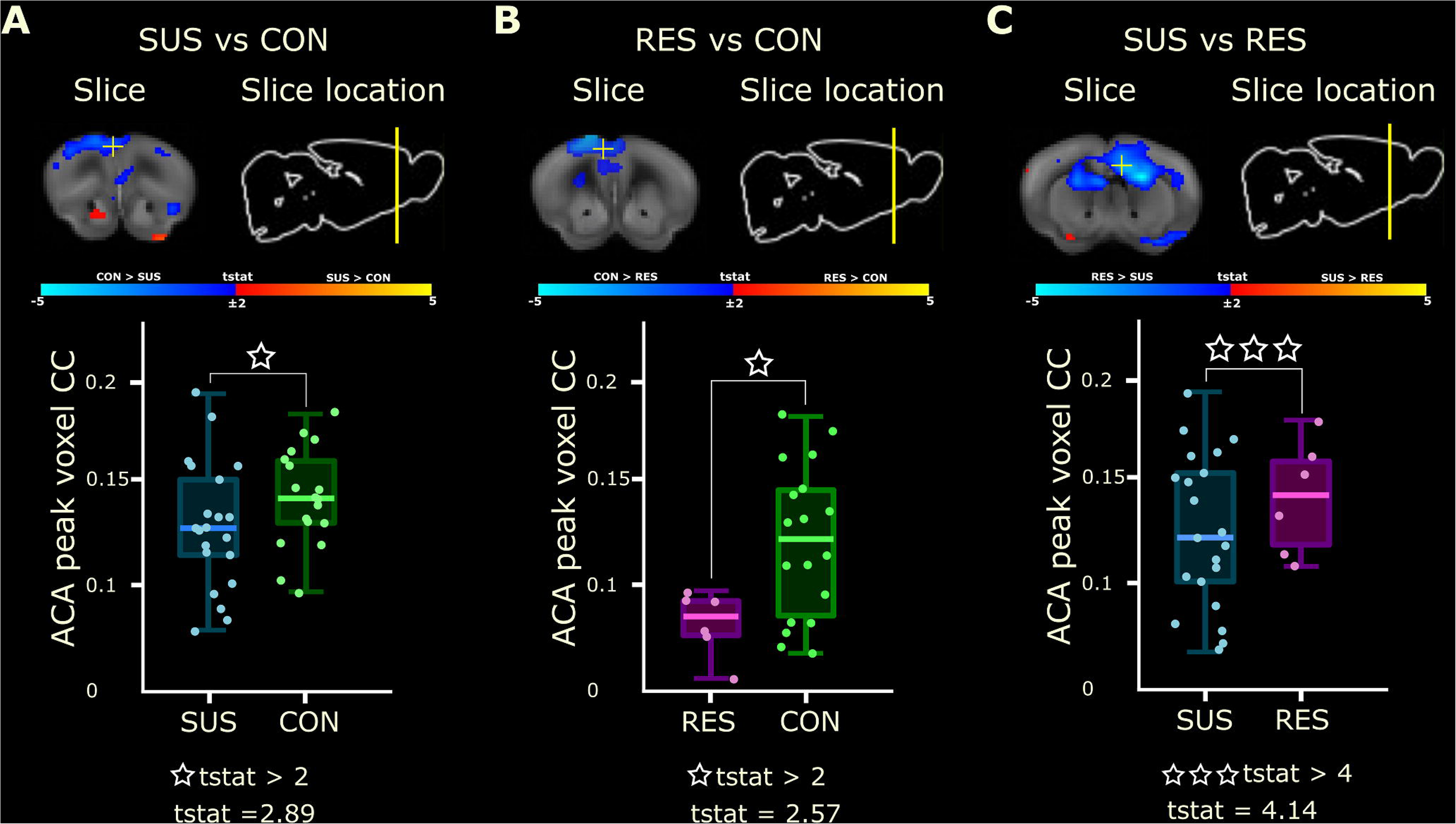
Group contrasts of vDG (top)—ACA peak voxel connectivity. (A-C) *Top panels*: Group contrast t-stat maps overlaid on Allen Mouse Brain Connectivity Atlas (2011) coronal slice illustrate vDG (top) seed-voxel connectivity and the anatomical location of the peak voxel (highest t-stat value) within the ACA cluster (white +) for SUS vs CON (A), RES vs CON (B), and SUS vs RES (C) mouse groups. Significant group differences are displayed as spectral intensity: red-yellow (t>2, cc=0.05, FWER-corrected) and blue-light-blue (t<-2, cc=0.05, FWER- corrected). *Bottom panels*: Subject-level box plots of vDG seed—ACA voxel correlation coefficients (CC) indicate a gradient effect of vDG–ACA connectivity that is generally stronger in undefeated CON mice compared to SUS (A, t(37)=2.89, p=0.006, FDR cut-off 0.05) or RES (B, t(22)=2.57, p=0.02, FDR cut-off 0.05) mice groups but is robustly increased in RES mice when compared to SUS animals (C, t(25)=4.14, p=0.0003, FDR cut-off 0.05). *Abv*: ACA, anterior cingulate area; RES, resilient; SUS, susceptible; vDG, ventral dentate gyrus; CON, control.

### Quantification of vDG seed-voxel findings with selected brain regions

We then sought to determine the magnitude of connectivity clustering between the vDG (top) source seed and five target brain regions; including, the anterior cingulate cortex, nucleus accumbens, basolateral amygdala, ventral tegmental area, and periaqueductal grey (**Fig. 6A**). Using group contrast t-stat maps we quantified the number and percent (relative to total seed region size) of voxels from region-specific connectivity clusters that showed a significant FC difference with the vDG (top) seed. The magnitude of FC was represented by the line thickness/colour bar between the vDG (top) seed and each target seed region (**Fig. 6B-D**). The magnitude of the FC was significant and most robust between the vDG and nucleus accumbens as well as periaqueductal grey regions in SUS compared to CON mice (**Fig. 6B**). The magnitude of the FC difference was also significantly higher in SUS compared to RES mice between the vDG and several limbic structures including the basolateral amygdala, ventral tegmental area, and periaqueductal grey (**Fig.6D**) supporting the previous analyses indicating hyperconnectivity between the vDG and other limbic regions within the SUS mouse brain. Quantification analyses further indicated a significantly increased magnitude of FC between the vDG and anterior cingulate cortex in RES compared to SUS mice (**Fig.6D**). These quantification analyses are supportive of the previous seed-seed and seed-voxel findings and highlight an important biological mechanism for resilience following exposure to CSDS.

**Fig. 6.**
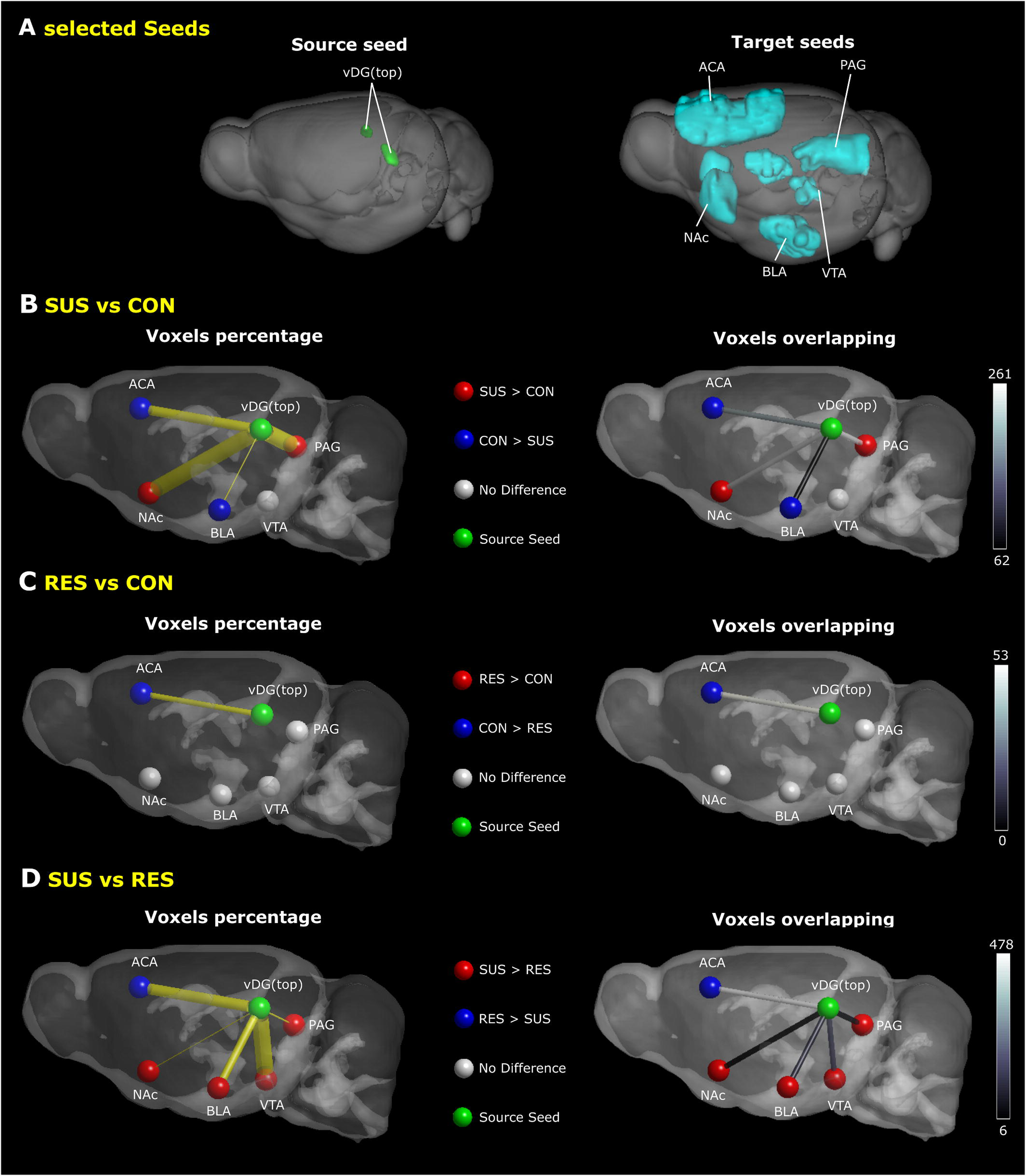
Quantification of vDG (top) seed-voxel functional connectivity (FC) with selected brain regions. A) 3D representations of a manually generated vDG (top) source seed (**A-left**) and 5 selected target regions (**A-right**) overlaid on Allen Mouse Brain Connectivity Atlas (2011) whole brain images. **B-D**) Schematic illustrations outline a quantification of significant vDG seed-voxel FC from Fig. 4 with selected target regions between SUS vs CON (**B**), RES vs CON (**C**), and SUS vs RES (**D**) mouse groups. SUS mice show preferential FC between the vDG and subcortical regions with the strongest change between the vDG–VTA regions when compared to RES mice (41.79%). Prominent vDG FC with the ACA region is favoured in CON mice compared to both RES (3.03%) and SUS (9.89%) mouse groups; however, RES mice show the largest vDG–ACA FC change (27.32%) relative to SUS animals. *Left panels* indicate the percent of voxels within a specified target region that showed a significant FC group difference with the vDG source seed. The magnitude of this percentage is represented by the line thickness (yellow connectivity bar). *Right panels* indicate the number of voxels within a specified target region that show a significant change in vDG FC. The black-white colour scaling bar (right) indicates the total number of significant voxels. Green-coloured spheres indicate the anatomical location of the vDG region. The direction of group differences between the vDG and target regions is represented by a red or blue coloured sphere. Regions with no group difference are indicated by a grey-coloured sphere. *Abv:* ACA, anterior cingulate area; BLA, basolateral amygdala; CON, control; NAc, nucleus accumbens; PAG, periaqueductal gray area; RES, resilient; SUS, susceptible; vDG, ventral dentate gyrus; VTA, ventral tegmental area.

## DISCUSSION

### General summary

CSDS is a well-validated model system for studies of the molecular mechanisms underlying individual differences in stress reactivity (18, 19, 22, 24, 49). The relevance of the model, particularly for the study of resilient versus susceptible animals, is based on the findings that 1) stress acts to precipitate MDD amongst a segment of the population and that 2) individual differences in stress reactivity moderate the impact of the stress on the risk for subsequent MDD. An understanding of the neurobiological basis for individual differences in susceptibility to chronic stress is thus of importance for an understanding of the pathophysiology of mood disorders

We sought to provide a comprehensive analysis of differences in neural connectivity that distinguish animals resilient or susceptible to the effects of chronic stress. We examined resting-state brain connectivity in SUS compared with RES mice focusing on the hippocampal vDG, a region critical for the expression of behavioral effects of CSDS (31, 32, 34, 36, 37, 52). The findings from the current *a priori* seed-seed and seed-voxel FC analyses distinguished SUS and RES phenotypes following CSDS. Thus, a variety of analytical approaches to the rs-fMRI data consistently showed enhanced FC of vDG with limbic regions critical for the activation of behavioral responses to chronic stress in SUS mice. In contrast, these same analyses revealed stronger FC of the vDG with prefrontal regions, including the anterior cingulate, a region known to moderate responses to stress in humans.

### Functional connectivity of the vDG in stress-susceptible animals

The rodent nucleus accumbens receives glutamatergic projections from the ventral hippocampus, which represents a primary input to the shell region (53–55). Our specific interest in the vDG as a seed region was based on evidence of relevant connectivity with stress-related brain regions(40) as well as its essential role in emotional regulation (38) stress susceptibility, depression, and antidepressant effects (3, 33, 37, 56–59). Of particular importance was the finding, across multiple analyses, of vDG hyperconnectivity with the nucleus accumbens in SUS animals. These findings are consistent with the wealth of studies showing that vDG projections to the nucleus accumbens are critical for the expression of depression- and anxiety-like behaviors following CSDS (33, 34, 36, 37, 60). SUS mice exhibit greater glutamatergic tone between the nucleus accumbens and ventral hippocampus compared to RES mice (34). Furthermore, excitation of the vDG increases susceptibility to social defeat in mice (33). Importantly, Muir et al (2020) showed that activity along the vDG – nucleus accumbens path was predictive of susceptibility to chronic variable stress in both male and female mice (60). These findings suggest that 1) the importance of the vDG – nucleus accumbens connectivity for stress susceptibility is not unique to the CSDS model and 2) that this pathway operates to define stress susceptibility in both males and females.

Our finding of increased FC between the vDG and ventral tegmental area in SUS mice is consistent with the role of the mesolimbic dopamine circuit in stress susceptibility and depression (61). Variation in dopamine neuron burst firing associates with susceptibility to stress in mice (18, 42, 62–64). In the CSDS model, RES and SUS mice use different behavioural strategies and reveal distinct activity patterns in dopamine terminals in the nucleus accumbens (41). It is interesting to note that the VTA and dopamine neuronal activity is influenced by communication between the vHPC and the nucleus accumbens (65).

The hyperconnectivity of the vDG with limbic regions in SUS animals included the basolateral amygdala. Hultman et al. (2018) used machine learning modeling of electrophysiological recordings to identify a neural circuit that predicted depressive-like behavior following CSDS (31). Increased activity within this network distinguished SUS animals. CSDS-related network activity was relayed through the amygdala and ventral tegmental area to converge in ventral hippocampus, suggesting greater amygdala - vDG connectivity associates with susceptibility. Chronic restraint stress in adult mice is also associated with greater amygdala connectivity to the ventral hippocampus (66, 67). There is an analogy with human clinical studies. An extensive meta-analysis of neuroimaging studies showed hyperconnectivity between the amygdala and the right hippocampus among MDD patients compared with healthy controls (68). The activation of stress responses is highly dependent upon activity in the right hemisphere, including the right hippocampus (69). Moreover, early life stress, which associates with greater susceptibility to chronic stress in later life, strengthens connectivity between the amygdala and hippocampus (70).

### Functional connectivity of the vDG in stress-resilient animals

We identified significantly greater FC between the vDG and prefrontal regions, notably the anterior cingulate cortex, in RES compared with SUS animals. There are important caveats concerning the alignment of the rodent and human prefrontal regions (71, 72). We assume that the mouse prelimbic and anterior cingulate regions bear homology to the human anterior cingulate (71), a region strongly implicated in MDD (45, 73, 74). Reduced connectivity of the prefrontal with the hippocampus is reported in patients with MDD (75, 76). Studies with animal models show that experimental procedures that enhance connectivity between the ventral hippocampus and prefrontal cortex produce antidepressant effects (36, 77). Activity within the ventral hippocampus – medial prefrontal cortex pathway is essential for the antidepressant response to ketamine (77). These findings are consistent with previous reports showing that hypoactivity in the mouse medial prefrontal cortex was associated with greater susceptibility to CSDS (78, 79).

The anterior cingulate cortex is associated with MDD in structural MRI studies (80) and essential for emotional and cognitive functions disrupted in MDD (45, 81–84). Alterations in anterior cingulate function and connectivity is proposed as a mechanism that manifests in features of MDD such as rumination, negative attentional bias, and emotion dysregulation that are characteristic features of depression. Moreover, anterior cingulate activity predicts treatment outcomes among depressed patients (85), a finding confirmed in an extensive meta-analysis (45). While connectivity of the anterior cingulate cortex to limbic regions is implicated in the emotional dysfunction characteristic of MDD, the precise role of altered connection with the ventral hippocampus remains to be clearly defined, although both regions are involved in emotional processing. One possibility is that increased connectivity between the ventral hippocampus and anterior cingulate may underlie improved emotional regulation associated with resilience.

The RES mouse brain was characterized by functional hyperconnectivity between subcortical (vDG or nucleus accumbens) and cortical regions including both the anterior cingulate and infralimbic prefrontal regions. There is reduced FC between the nucleus accumbens and connecting reward, executive, default mode, and salience networks in MDD patients (86) FC strength between the nucleus accumbens shell region and the subgenual/pregenual anterior cingulate cortex associates with anhedonia in MDD patients (87). Enhanced activity between the infralimbic prefrontal cortex and nucleus accumbens regions may be a mechanism that supports resilience following exposure to CSDS. Indeed, direct glutamatergic projections exist between the medial prefrontal cortex and the nucleus accumbens in rodents (88, 89) and stimulation of the medial prefrontal region elicits increased extracellular glutamate levels in the nucleus accumbens (90). Acute optogenetic stimulation of medial prefrontal afferents in the nucleus accumbens reduced social avoidance during social interaction testing in CSDS mice (34); an effect that further defined previous work of increased medial prefrontal activity as pro-resilient (78).

Together, these findings support a consensus view that enhanced connectivity with prefrontal regions is protective and that effective coordination between this area and at least some connections, specifically hippocampal and nucleus accumbens regions, represents a tractable and translatable mechanism underlying a resilience to the developments of depression-related pathophysiology.

### Limitations

The CSDS paradigm has emerged as an impressive model for the study of transcriptomic and behavioral mechanism of depression. Transcriptional profiles resulting from exposure to CSDS show highly significant overlap with associated with MDD in post-mortem human brain (24). However, the model remains less appropriate for females. Alternative paradigms have been implemented in females with comparable depressive-like effects to those obtained by CSDS in males (92); however, comparing sex effects between different models remains challenging to interpret (93). Stressors, particularly social stress, may have very different functional significance for males and females even with the same experimental protocols. We acknowledge the exclusive use of male animals as a limitation and suggest these findings are considered preliminary requiring replication and extension with females using appropriate chronic stress protocols. Additionally, it is important to replicate these findings using alternative chronic stress models to ensure comparable results across various models of resilience.

### Conclusions

We found highly unique patters of FC between animals defined as susceptible or resilient using a well-established rodent model. The enhanced connectivity in SUS animals of the ventral hippocampus to limbic regions implicated in the activation of stress is consistent with existing literature in the study of chronic stress. A highly novel finding here is that of the hyperconnectivity of ventral hippocampus to prefrontal cortex as a consistent feature of RES animals. Neuroimaging analyses of clinically-relevant animal models thus allow for identification of novel pathways underlying individual differences in responses to chronic stress. An advantage of the neuroimaging approach is that it is operationally and analytically comparable across rodents and humans, thus providing an important bridge to human neuroscience and the study of the neural basis for psychiatric disorders.

## Acknowledgements

Funding for this project was provided by the Hope for Depression Research Foundation (HDRF) (MJM).

## Supplementary Information

**Supplemental Figure S1. Behavioural characterization of control, susceptible and resilient mice.** Subject-level social interaction (SI) ratios (time spent in the interaction zone with social target present/time spent in this zone with the target absent) from identified mouse groups (mean ± SEM). SI testing characterized defeated animals as ‘susceptible’ (SUS, n=20) or ‘resilient’ (RES, n=6) with an SI ratio less or greater than 1, respectively. Undefeated control (CON, n=18) mice were also subjected to SI testing. SUS mice scored significantly lower in SI testing compared to both CON (***p<0.0001) and RES (**p<0.001) mouse groups. *Abv*: CON, control; RES, resilient; SUS, susceptible.

