## Supplementary Information - Supplementary Figure 1 for "Resting-state fMRI reveals altered functional connectivity associated with resilience and susceptibility to chronic social defeat stress in mouse brain"

Affiliations:

Running title: Functional connectivity and social defeat in mice

**Contents**

The following comprises Supplementary Figures that were not included in the printed version due to space constraints. These data detail:

**Figure S1**: Social interaction (SI) test data

**Supplementary Results**

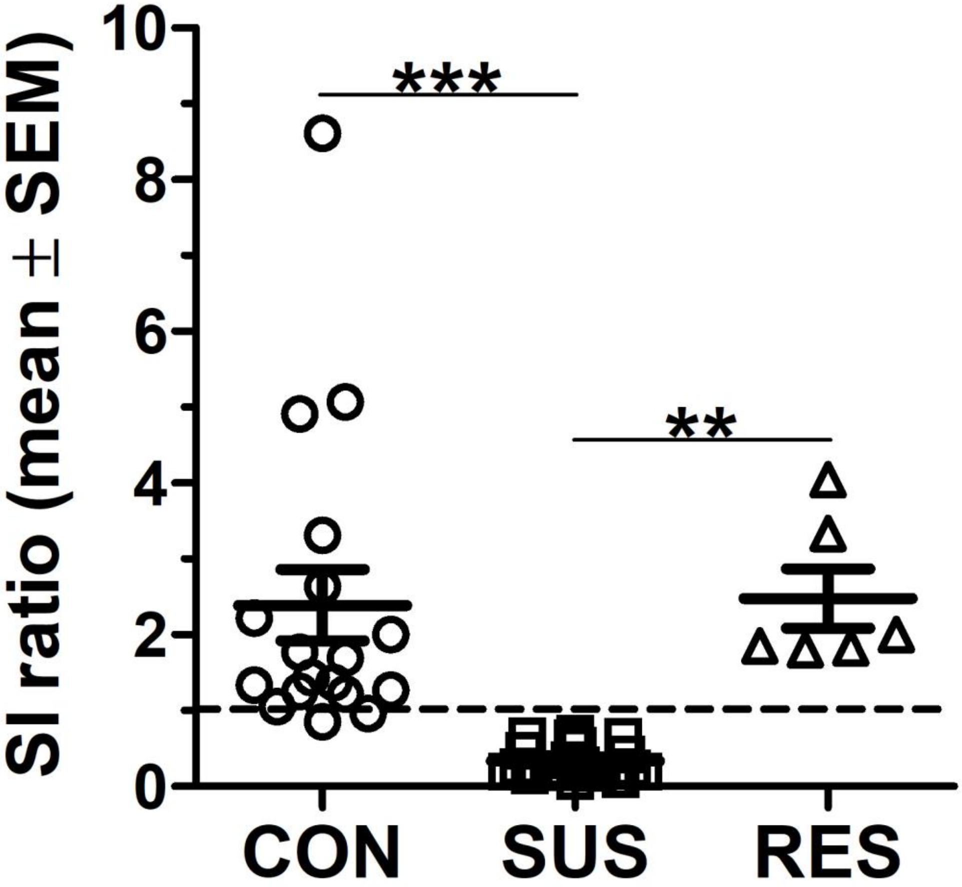


**Supplementary Fig. S1. Behavioural characterization of control, susceptible, and resilient mice**

Subject-level social interaction (SI) ratios (time spent in the interaction zone with social target present/time spent in this zone with the target absent) from identified mouse groups (mean ± SEM). SI testing characterized defeated animals as 'susceptible' (SUS, n=20) or 'resilient' (RES, n=6) with an SI ratio less or greater than 1, respectively. Undefeated control (CON, n=18) mice were also subjected to SI testing. SUS mice scored significantly lower in SI testing compared to both CON (***p<0.0001) and RES (**p<0.001) mouse groups. *Abv*: CON, control; RES, resilient; SUS, susceptible.
